# HiGlass: Web-based Visual Exploration and Analysis of Genome Interaction Maps

**DOI:** 10.1101/121889

**Authors:** Peter Kerpedjiev, Nezar Abdennur, Fritz Lekschas, Chuck McCallum, Kasper Dinkla, Hendrik Strobelt, Jacob M Luber, Scott Ouellette, Alaleh Azhir, Nikhil Kumar, Jeewon Hwang, Soohyun Lee, Burak H Alver, Hanspeter Pfister, Leonid A Mirny, Peter J Park, Nils Gehlenborg

## Abstract

We present HiGlass, an open source visualization tool built on web technologies that provides a rich interface for rapid, multiplex, and multiscale navigation of 2D genomic maps alongside 1D genomic tracks, allowing users to combine various data types, synchronize multiple visualization modalities, and share fully customizable views with others. We demonstrate its utility in exploring different experimental conditions, comparing the results of analyses, and creating interactive snapshots to share with collaborators and the broader public. HiGlass is accessible online at http://higlass.io and is also available as a containerized application that can be run on any platform.

## Background

The development of chromosome capture assays measuring the spatial contacts between two or more regions of the genome is essential for elucidating how the structure and dynamics of the genome affect gene regulation and cellular function [1,2]. Genome-wide maps of chromosomal interactions obtained by techniques such as Hi-C have revealed features of genome organization such as compartmentalization, i.e. spatial segregation of active and inactive regions of the genome, topologically associating domains (TADs), and associated peaks of contact frequency (often referred to as loops) [1,3–5]. Hi-C maps have helped implicate changes in genome organization in a variety of disorders, including acute lymphoblastic leukemia [6], colorectal cancer [7], and limb development disorders [8]. More fundamentally, they provide insights into the mechanisms by which genome conformation structures arise, are maintained, and change over time [9–11]. Major efforts like the 4D Nucleome Network and the ENCODE project are generating such data at large scale across different cell lines and conditions with the aim of understanding the mechanisms that govern processes such as gene regulation and DNA replication as well as to cross-validate the results from different experimental assays [12,13].

Despite the large amounts of generated Hi-C data, major challenges remain in (i) identifying known features unambiguously [14]; (ii) discovering new features; (iii) establishing relationships between Hi-C features and known (epi)genetic profiles; (iv) establishing the effects of various genetic, biochemical and physical perturbations on chromatin organization, assessing meaningful differences between cell types [15], and changes across the cell cycle and along differentiation pathways [16]. These challenges necessitate the development of methods to visually explore, compare and share not only the raw data, but also related datasets and derived analysis results. An effective visualization platform needs to meet the following criteria: (1) Provide researchers with the means to explore their data and look for patterns that may help to interpret the results of experiments and generate hypotheses. (2) Enable efficient comparison by juxtaposition or other means of different samples or conditions and integration of both similar and heterogeneous data types. (3) Allow researchers to overlay computationally derived annotations to visually validate analytical results as well as to compare the outputs of different data processing pipelines. (4) Enable sharing of results with collaborators and the public. And crucially, an effective platform does this all in a fast, intuitive, and accessible manner.

To obtain genome conformation capture maps, raw Hi-C sequencing data are processed to identify proximity ligation events representing captured contacts between genomic loci, which are then binned to form contact matrices [17–19]; see Lajoie et al. [20] and Ay & Noble [21] for reviews of Hi-C data processing. The discovery and elucidation of genome organizational principles and mechanisms, however, also requires sophisticated visual tools for exploring features relevant at scales ranging from tens to millions of base pairs [18,22,23]. Given the multiscale features of genome organization, it is crucial that such visualization tools support comparison across multiple scales and conditions as well as integration with additional genomic and epigenomic data. Existing tools provide different ways of displaying contact frequencies, such as rectangular heatmaps, triangular heatmaps, arc plots, or circular plots, and different degrees of interactivity ranging from static plotting to interactive zooming and panning, as well as different degrees of integration with other genomic data types [18,24–29]. While tools such as Juicebox [18] and Genome Contact Map Explorer [30] provide synchronized exploration of multiple contact maps, they lack an interface for dynamically arranging the views of several Hi-C datasets, and customizing the levels of synchronization between loci, zoom levels, and samples. Furthermore, none provide an interface for continuous panning and zooming of the sort popularized by web based geographical and road maps.

To address these shortcomings, we created HiGlass, an open source, web-based application designed to support multiscale contact map and genomic data track visualization across multiple resolutions, loci, and conditions (http://higlass.io, Supplementary Methods). HiGlass was built with an emphasis on usability. It provides an interface for continuous panning and zooming across genome-wide data. To facilitate comparison and exploration, HiGlass introduces the concept of *composable linked views* for genomic data visualization (Fig. 1). Each *view* in HiGlass is a collection of 1D and 2D tracks sharing common genomic axes. Views can be filled with data tracks, resized, arranged spatially, and linked to synchronize their axes by location or zoom level. This approach enables users to interactively compose the layout, content, and synchronization of locus, zoom-level, and other properties across multiple views (Fig. 1). By creating, sizing, arranging, and linking individual views, users can create custom compositions ranging from the juxtaposition of two or more heatmaps to sophisticated arrangements of views containing matrices, tracks, and *viewport projections* mapping the extents of one view inside another (Fig. 1, 2, Supp. Fig. 1). We demonstrate how HiGlass has been used to detect and analyze novel features in Hi-C data, and to visualize, validate, and compare tools for detection of known features.

**Figure 1.**
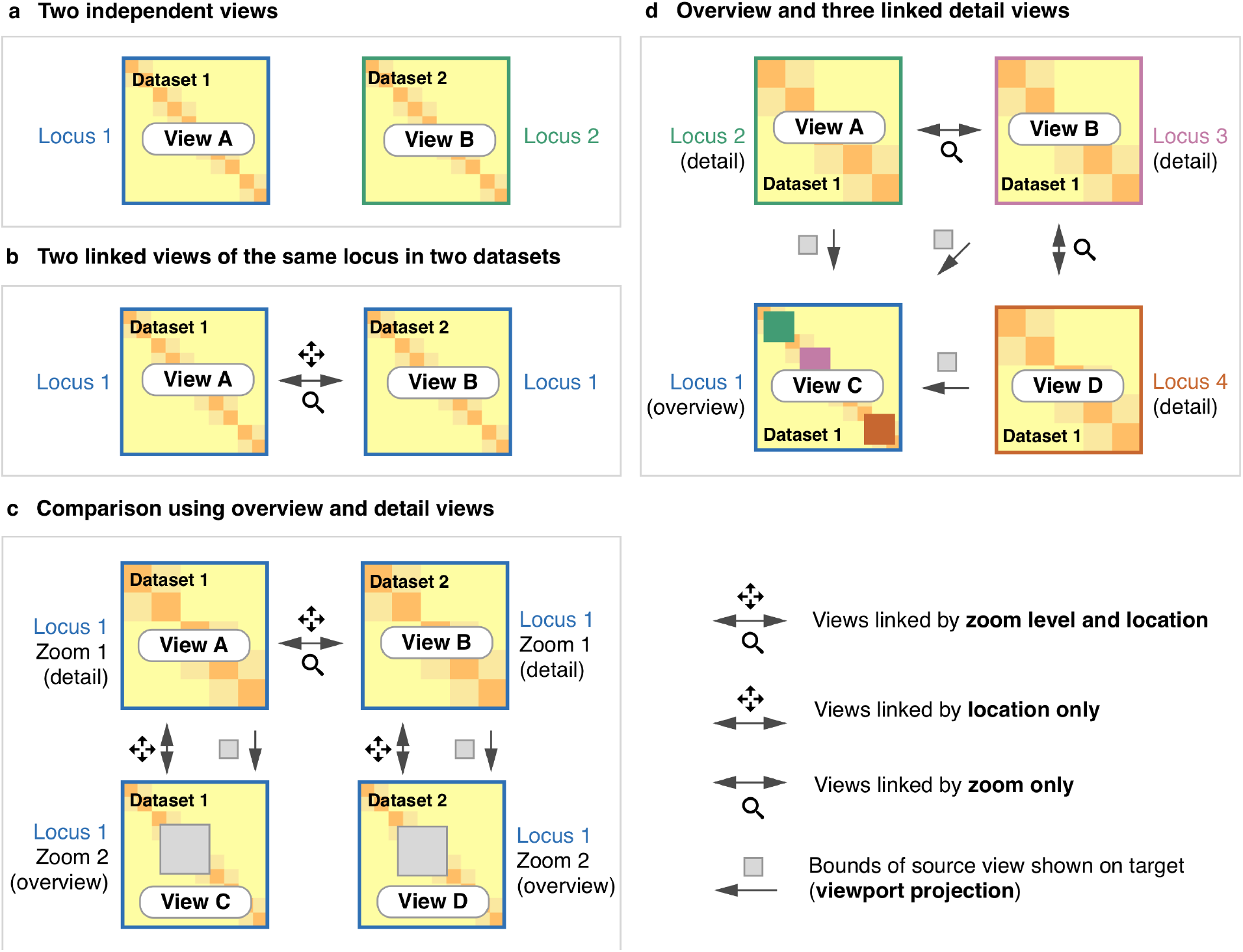
Schematics of different ways that views can be linked. Multiple views of the same (b) or different datasets (a,b,c) can be composed and linked to facilitate data exploration and comparison. Two independent views of different samples provide free and independent exploration of each sample (a). Linking by zoom and location enforces the same scale and location in both samples (b). Zoom linking maintains the same scale while allowing free independent manipulation of the location (d). By linking location and leaving zooming free, one set of views can show an overview of a high resolution region (c). Displaying the extent of one view in another is referred to as a viewport projection in this manuscript and shows where a detail view is located in an overview (c,d). The process of linking views is illustrated in Figure 6.

**Figure 2.**
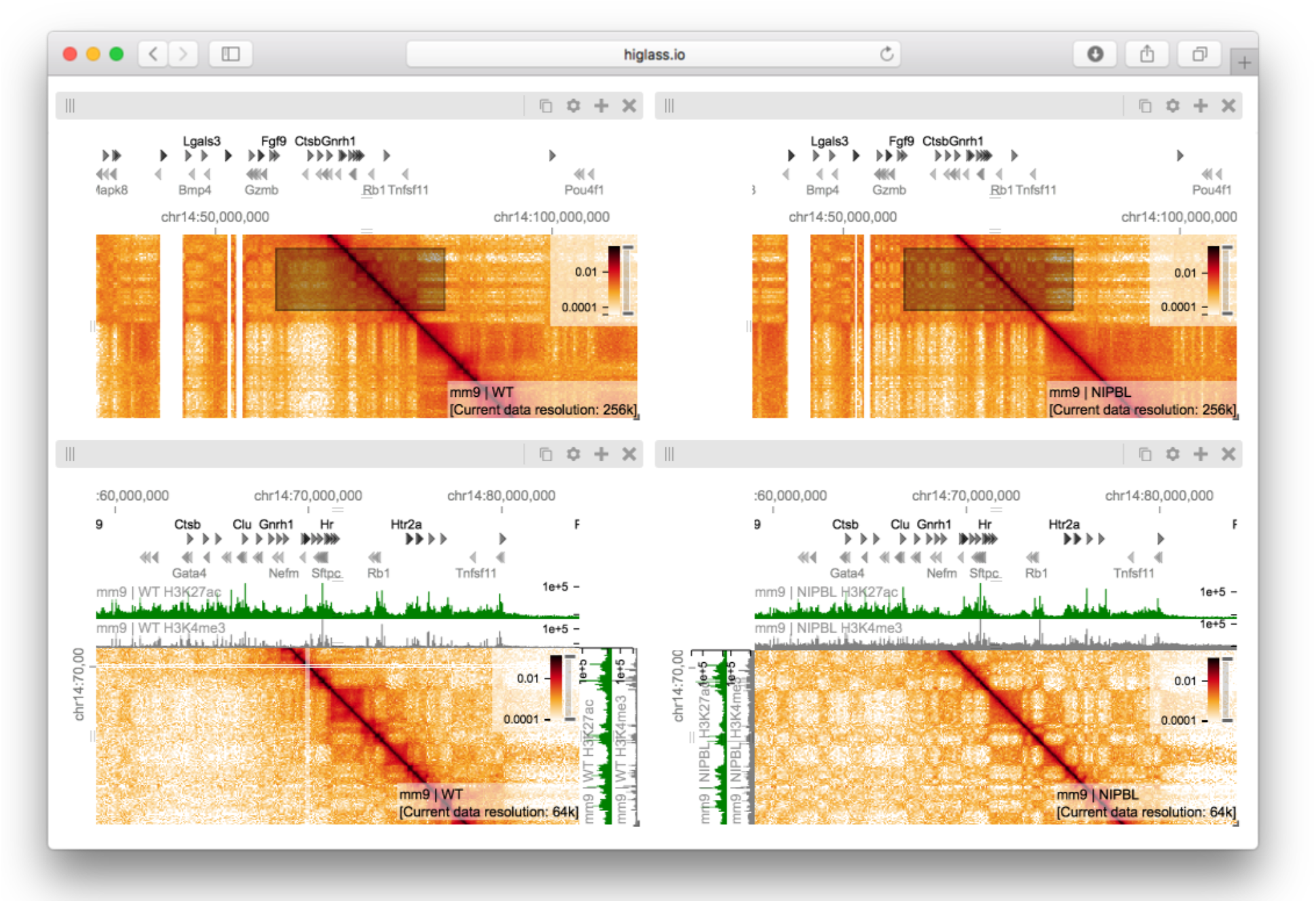
A view composition highlighting the results from Schwarzer et al. with data from WT (left), and mutant (ΔNipbl, right) samples [33]. The top two views are linked to each other by zoom and location such that they always display the same region at the same resolution. Comparing the control (left) and mutant (right) condition at this zoom level reveals the bleaching of TADs in the gene-poor region in the lower right hand part of the maps. The bottom two views, also linked to each other by zoom and location, show a zoomed in perspective where a more fragmented compartmentalization of the ΔNipbl mutant (right) as compared to WT (left) can be seen. The black rectangles in the top views, which are referred to as viewport projections in HiGlass, show the positions and extent of the bottom views (Supp. Fig. 2). The white lines in the bottom left panel are a result of bins filtered during matrix balancing. An interactive version of this figure is available at http://higlass.io/app/?config=Tf2-ublRTey9hiBKMlgzwg.

Multiple views within the same browser window, with synchronized panning and zooming, allow fast comparison of Hi-C maps for different samples/conditions. Views can, in the simplest case, be arranged to show the same location at the same zoom level across multiple samples (Fig. 4 and Fig. 5). In other cases, the investigator may wish to view multiple loci within the same sample (Fig. 1b and Supp. Fig. 4). More complex arrangements can pair views with differential zoom levels in a context-detail arrangement (Fig. 2 and Fig. 3) [31]. View compositions serve to display data at multiple scales, to corroborate observations with other types of evidence and to facilitate comparisons between experiments. As a web-based tool, HiGlass also supports storing and sharing of view compositions with other investigators and the public via hyperlinks. The tool can be used to access selected public datasets at http://higlass.io or it may be run locally and populated with private data using a provided Docker container. It can also be embedded within other applications to provide a component for displaying Hi-C or other genomic data [32].

**Figure 3.**
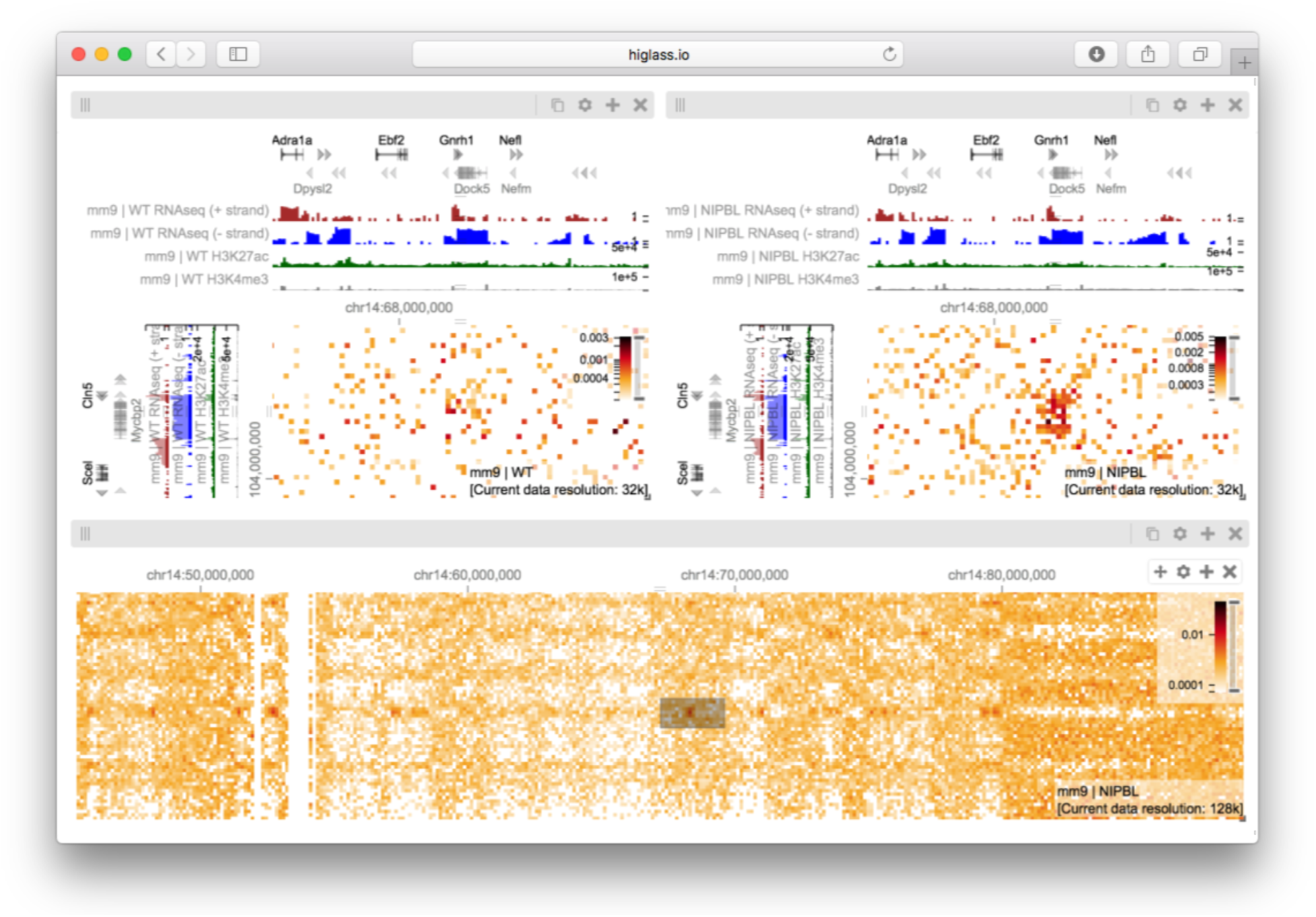
A view composition containing two views linked by location and zoom (top) and an independent (unlinked) zoomed out overview (bottom) (Supp. Fig. 3). The two views on top show data from chromosome 14 (mm9) in the wild type and ΔNipbl conditions, respectively. The bottom view shows data from the mutant condition as well as a projection of the viewport visible in the top views. The patch visible in the ΔNipbl condition (top left) is notably absent from the control (right). The gene annotations, RNAseq, H3K27me and H3K4me3 tracks show the presence and transcription of the *Dock5* and *Mycbp2* genes on the - strand as well as the presence and transcription of the *Gnrh1* and *Cln5* genes on the + strand. An interactive version of this figure is available at http://higlass.io/app/?config=Q5LdNchQRLSZOyKsTEoiw.

## Results

### Exploring and comparing different experimental conditions

To illustrate the utility of composable linked views in exploring different experimental conditions, we used HiGlass to highlight key results of a recent study showing the effect of induced deletion of the cohesin loading factor *Nibpl* on chromosome organization in adult mouse hepatocytes [34]. We obtained Hi-C contact data and binned it at multiple resolutions starting at 1 kb for wild type (WT) and ΔNipbl primary hepatocytes (Supp. Methods). We loaded both samples as separate views (Fig. 2, top) and linked them via location and zoom level. With the two linked views, we could navigate to regions clearly showing the disappearance of features in the ΔNipbl condition. We also added views of genomic positions and locations of individual genes that move in sync with Hi-C maps, allowing to examine how changes in Hi-C in different genic contexts. For example, in the gene-poor region from chr14:80 Mb to chr14:100 Mb of mm9, we observe a robust loss of near-diagonal contact enrichment patterns. We identify the contact patterns that disappeared as TADs in a strict sense because they do not show the long range associative “checkerboard” pattern of A/B compartmentalization, a feature that remains intact in the ΔNipbl condition [35]. In contrast, in the relatively gene-rich region upstream of chr14:80 Mb, we see an enhancement of the checkered pattern and the emergence of a finer division of A/B regions in the ΔNipbl condition. To explore this region more closely, we created two additional linked views for WT and ΔNipbl and navigated to the region between chr14:50 Mb and chr14:70 Mb (Fig. 2, bottom). Adding H3K4me3 and H3K27ac ChIP-seq signal tracks revealed that these marks, while similar between conditions, correlate more strongly with the compartmentalization pattern in ΔNipbl. Finally, we used a viewport projection to mark the position of the bottom views relative to the top, resulting in the complete view composition shown in Fig. 2. This interactive visual recapitulation of key results from Schwarzer et al illustrates how synchronized navigation across loci and resolutions by linking views between multiple conditions facilitates the exploration of the complex effects of global perturbations on chromosome organization at multiple scales.

Using the same view composition we noticed the appearance of a new feature, small dark patches (“blotches”) away from the diagonal in the ΔNipbl condition. To investigate these patches we created a new composition containing an overview and two zoom- and location-linked detail views (Fig. 3). By using the overview to find patches and comparing them using the detail views, we established that they are more enriched in the mutant condition than in the wild type, that they represent strengthened interactions between pairs of short active regions (type A compartment) and that they tend to be aligned with annotations of long multi-exonic genes. Including RNA-seq and ChIP-seq tracks let us see that the genes which align with these patches are virtually always transcriptionally active. These observations are reminiscent of a recent ultra high resolution Hi-C study in mouse ES and neural cells, where the long-range contact enrichment between pairs of expressed genes was found to correlate with both expression level and the number of exons, and agrees with similar strengthened patterns observed after degradation of cohesin in a human cell line [36,37]. Not only do composable linked views provide convincing support that the absence of cohesin loading leads to strengthening of global genome compartmentalization, but they also hint that, at finer scales, long range and inter-chromosomal contact enrichment and its response to cohesin loss are influenced by transcriptional parameters such as expression output and splicing activity.

### Comparing the results of feature callers

Analysis of genomic data usually involves identification and annotation of various “features” that range from calling sequence variants to detecting complex patterns of interactions in Hi-C maps. Often, the first step in characterizing the quality of a caller is a visual inspection to verify that the regions it annotates match the expectations of the human analysts. In the case of ChIP-seq data, for example, peak callers identify regions where proteins bind [38] and an analyst would verify that the regions contain an elevated number of read counts relative to the surrounding regions. In Hi-C data, topologically associated domain (TAD) callers identify regions of increased contact frequency in contiguous loci (e.g. along the diagonal in a Hi-C map) [3,4]. In contrast to 1D peak callers, TAD callers demarcate square regions of interest in a Hi-C map. This makes comparison more complicated as the results often need to be placed next to each other, rather than simply stacked on top of each other. Results from multiple callers run on multiple replicates further complicate the task of comparison.

To address the first issue of comparing feature calls on 2D maps, we obtained data for the comparison of seven algorithms that identify TADs from Forcato et al and created a view composition consisting of eight different views (Fig. 4) [14]. Seven views show called TADs overlayed on top of the same Hi-C map, with the eighth map showing map unobstructed by markers of called TADs. All views were then synchronized by zoom and location. By ensuring that each view always showed the same genomic region, we can compare the results at the same scale and location. Clearly visible in this comparison is the lack of consensus between the different available TAD callers. Few regions are consistently called by more than one caller. The lack of consensus is also evidenced by the variation in the size of the called TADs. While this variation in size is demonstrated empirically by Forcato et al., seeing the calls overlaid on the raw data can reveal that some are not only on the same scale as the larger compartment features, but also overlap with compartmental transitions (Supp. Fig. 5). Downstream analysis based on such TAD calls should therefore consider whether phenomena attributed to TADs can also be attributed to other features of Hi-C.

**Figure 4.**
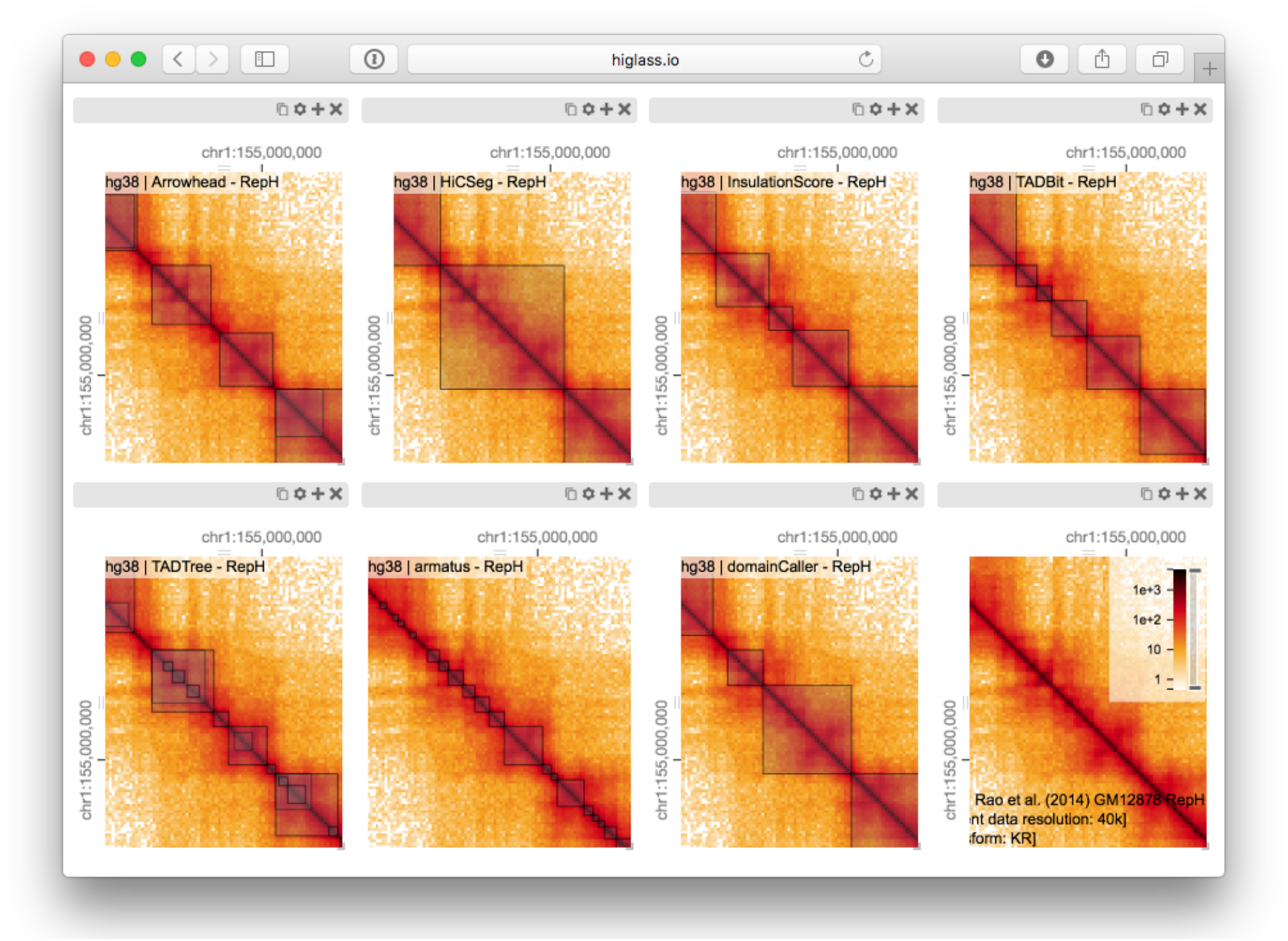
Eight views linked by location and zoom (Supp. Fig. 5). Each view shows the calls made by a single TAD caller overlaid on the matrix on which they were called. There is little consistency between the results of the different callers and large variation in the size of the TADs. The last view (bottom right) shows a matrix with no overlay. An interactive version of this figure is available at http://higlass.io/app/?config=IPCHmdOQR4CDY2sqj5VJHQ.

In addition to the differences between TAD calls among different callers, there are differences in the calls produced by a single TAD caller on different replicates. Such differences may be attributable to variations in signal-to-noise (e.g. quality and depths of different sequencing runs and differences in library complexity between replicates). Furthermore, by looking at the results of 7 different TAD callers among 10 experiments we can see that consistency within a caller does not imply consistency between callers. Such views also reveal more subtle differences. Some callers, for example, partition nearly all of the genome into a contiguous sequence of “TADs” (HiCseg [39], insulationScore [40] and TADbit [41]), while others (Arrowhead [5], TADTree [42], domainCaller [3] and Armatus [43]) call discontinuous intervals, and some methods allow for overlap and/or nesting [14]. Such differences raise meaningful questions about what data patterns are used to define TADs in different studies, how robustly different algorithms can capture any given pattern type, and how the findings from one study can be translated to those of another. These issues are further underlined by recent experimental perturbations of chromatin architectural factors, such as the *Nipbl* deletion study above, which reveal that segmental annotations based solely on local contact enrichment cannot all be attributed to the same organizational process inside the nucleus and that standard Hi-C maps reflect an interplay of distinct dynamic processes averaged over a cell population.

### Creating interactive snapshots of genome-wide data

In addition to exploration and interpretation, visualization is an essential tool for the communication of scientific findings. With the increasing use of high-throughput sequencing and genome-wide assays, screenshots of genome browsers have become common in computational genomics. Such figures convey the relationship between one (in the case of conventional genomic data) or two (in the case of chromosome conformation data) loci and some measure such as read coverage or fold change in coverage. In publications, the extents of these plots are limited by the space and resolution available on the printed page. This compels authors to show one or two loci that most clearly demonstrate the effect they are describing. The original data is archived in repositories such as the Gene Expression Omnibus (GEO). A user who wishes to explore additional examples or view the data using a different visual representation requires a non-trivial human effort to a) locate the data in the appropriate repository b) establish which files correspond to which figures and c) prepare, convert and load the data into a genome browser or viewer. This arrangement hinders communication, reproducibility and further analysis by dissociating the raw genome-wide data from the publication describing it.

With HiGlass, authors can produce links to interactive figures that can be shared with collaborators or the public. These links point to HiGlass view compositions that can show all of the genome-wide data used to produce a figure. These compositions are centered on one or more loci but can be navigated to other locations. Generating a link to a view composition stores all of the information necessary to reproduce it, including the data sources, track types, and synchronization links on the hosting server. This “view configuration” can also be stored as a file that can be shared with collaborators. Similar functionality was pioneered by the UCSC Genome Browser [44], where users could create “Track Hubs” hosting their own data and then share session links to genome browser views incorporating their data. Similarly, HiGlass users can run their own server locally and share links pointing to local data as well as data hosted on remote servers.

In contrast to most existing tools, HiGlass stores a declarative JSON representation of the current view configuration into its local database rather than the browser URL, which has a limited character length. HiGlass generates a link referencing the view configuration when the user selects to share their view composition. Without the need to encode every aspect of the visualization in the space-constrained URL, we can include more metadata about the how the tracks are styled and linked, the data sources and the synchronization options. This JSON state representation can either be saved locally or stored in HiGlass’s database and shared as a link to an interactive figure (Figures 2,3,4,5). By capturing the current composition and storing its complete state on the server, we create the opportunity to integrate HiGlass with tools for documenting and exploring the provenance of the composition to better understand the steps that the analyst took to reach their conclusions [45].

### Feature overview and comparison with other viewers

The major strengths of HiGlass are smooth navigation, multi-view comparison, comprehensive selection of track types, and containerized deployment. Of the existing browsers, only HiGlass and Genome Contact Map Explorer (GCME) provide a continuous interface for panning and zooming across loci and resolutions. Other tools, such as Juicebox, Juicebox.js [46], the Washington University Epigenome Browser (WUEB) and the 3D Genome Browser show data at fixed discrete zoom levels. To compare data, Juicebox, Juicebox.js, GCME and HiGlass offer the opportunity to place heatmaps side by side and navigate multiple Hi-C maps simultaneously. Of these, only HiGlass lets users select which heatmaps to synchronize or whether to synchronize by location, zoom, or both. This is critical for the creation of task-specific view compositions, for example to support overview and detail or multiple comparisons. Furthermore, no other tools let users establish connections between views (viewport projections) so as to display the location of one view within another (Fig. 2, Fig. 3).

The separation of data retrieval and rendering in HiGlass makes it easy to create new track types. HiGlass already supports horizontal triangular heatmaps (Fig. 5), vertical triangular heatmaps and 2D heatmaps (Fig. 2,3,4) for viewing Hi-C data as well as tracks for showing 2D annotations (Fig. 4 and Fig. 5). This is in contrast to other viewers such as Juicebox, Juicebox.js and GCME which display only 2D heatmaps or WUEB and the 3D Genome Browser which only display horizontal triangular heatmaps. Heatmaps in HiGlass are highly configurable. Color scales can be synchronized and tuned, and can be adjusted to display both linearly and logarithmically scaled data, an option also present only in GCME. Such features are crucial because the dynamic range of intra-chromosomal contact frequency spans several orders of magnitude. Genomic signal tracks can be displayed using lines, bars or points. Other track types such as gene annotations, rotated 2D annotations (Fig. 5) and generic 1D annotations are also directly supported. HiGlass supports selectable synchronized scaling between values in different tracks as well as the ability to fix heatmap color scales to a defined data range. It supports SVG export as well as JSON view configuration and link export for sharing (see Results section “Creating interactive snapshots of genome-wide data”).

**Figure 5.**
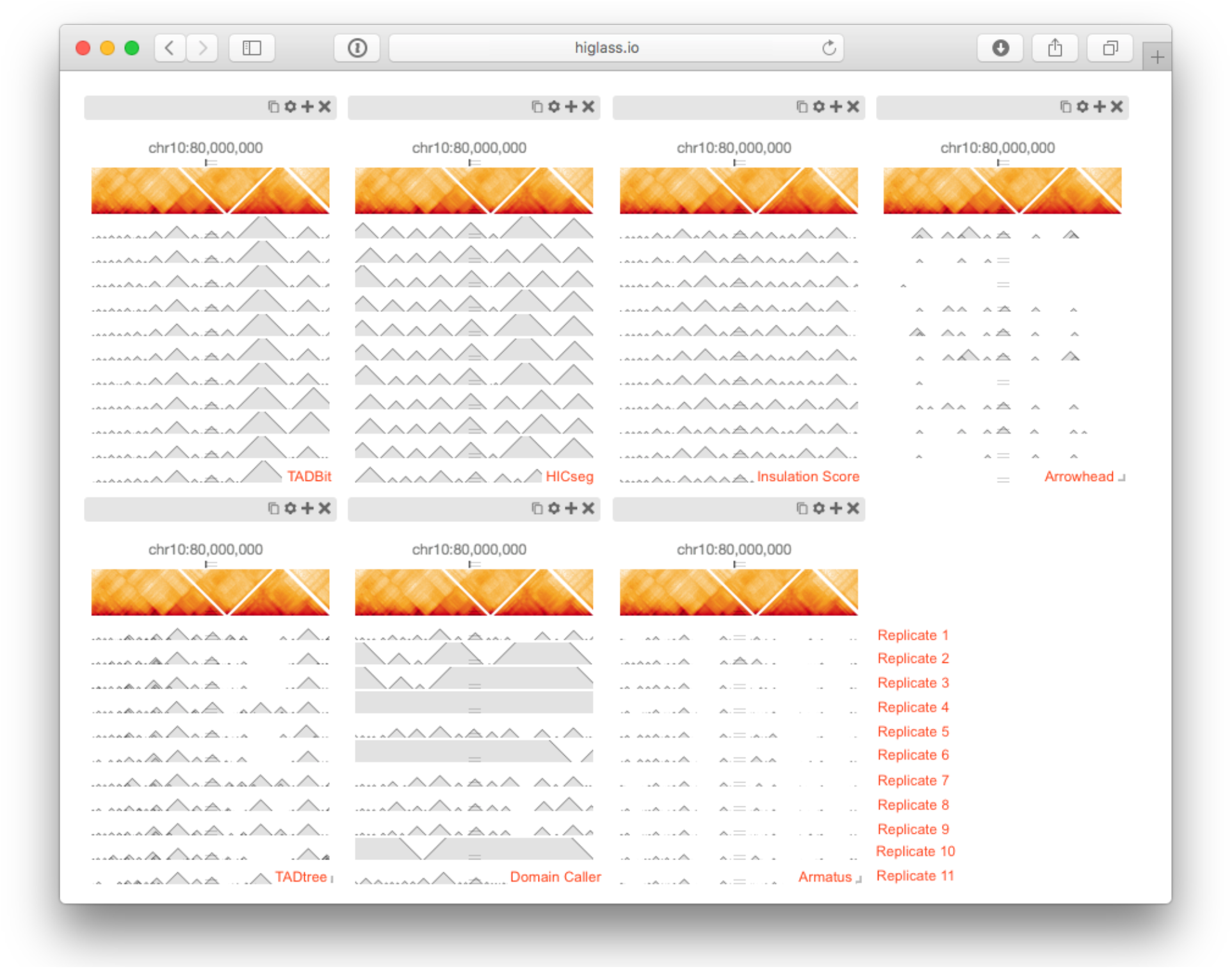
The seven views shown here show tracks in a horizontal configuration at the same location and zoom level (Supp. Fig. 6). Each view shows the output of a particular TAD caller (TADBit, HiCseg, Insulation Score, Arrowhead, TADtree, Domain Caller, and Armatus, from left to right, top to bottom). Each of the bottom tracks shows the output for a single replicate. The matrix on top contains data from a combination of all replicates. An interactive version of this figure is available at http://higlass.io/app/?config=JALHH-HzQGeJCaJaU9EwTA. The caller names and replicate labels were added for clarity.

For deployment, we provide a Docker container for HiGlass which can be run locally and populated with private or shared data (Fig. 6). This makes it possible for individuals to view local files or for laboratories to create instances shared within an internal network. Such instances can be used to isolate both data and shared interactive figures from the public. Laboratories can also set up public instances to share data and figures outside of the local network. The ability to set up public and private instances is also available for the Washington University Epigenome Browser but absent from other tools. Because Juicebox and Juicebox.js can load remote files, similar functionality can be approximated by controlling data access at its point of storage. Without a database, however, it is difficult to obtain lists of available track and their associated visual encodings from within the viewer itself. HiGlass makes it possible to not only collect sets of tracks locally but can connect to and obtain tracks from any number of different remote instances, such as the one at http://higlass.io.

**Figure 6.**
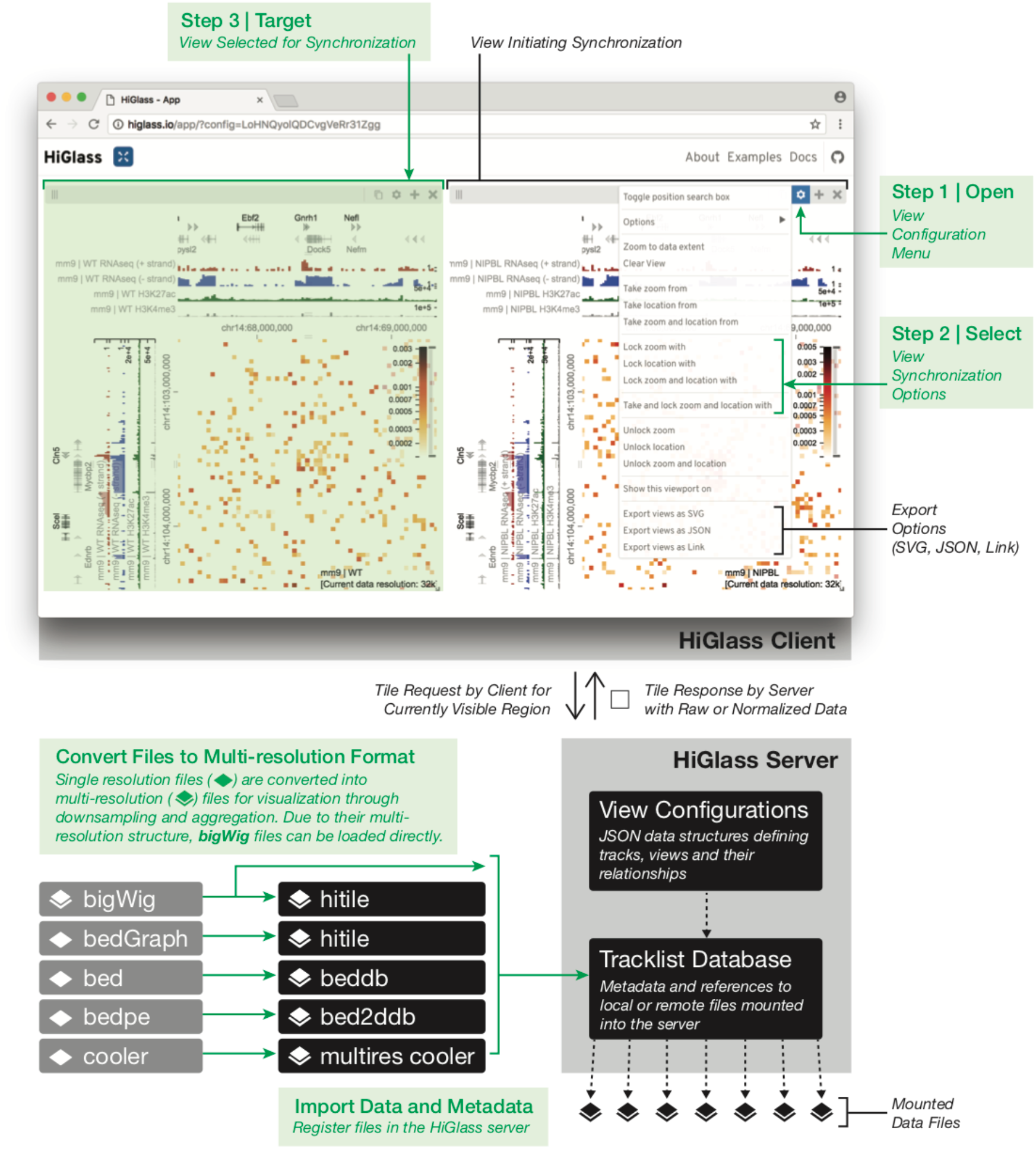
A schematic of the data flow and user interface of HiGlass. Starting from the bottom-left, single resolution formats used for genomics data are converted to their multi-resolution counterparts. Files in the bigWig format, which is a native multi-resolution format, are directly converted to the hitile format compatible with HiGlass. The multiresolution files are then loaded into the HiGlass server using a command line tool. The HiGlass client (top half) communicates with the server by issuing ‘tile requests’ for the data that is currently visible in the user’s browser. The server responds with raw data which the client renders into vertical, horizontal and 2D tracks. Within the client, users can zoom and pan around the data or select views with which to synchronize the location, zoom level or both. View synchronization is initiated from one view and tied to another.

## Conclusions

Using HiGlass to create the linked views shown in Figures 2 and 3 enabled us to interactively explore the data generated by Schwarzer et al. across different conditions, zoom levels and loci [47]. This gave us not only a clearer understanding of the results but also the ability to see them in a genic context, but also allowed us to find unexpected patterns, relate them to histone patterns and gene expression, and rapidly gather observations to be used in generating new hypotheses. We used a different composition of views to show and compare the results of seven different TAD callers in a single window [14]. This let us compare the variation among different TAD callers, and of the same caller, across different replicates (Fig. 5), as well as with the original data that the calls were generated from (Fig. 4). These figures highlighted the inconsistency in the results between separate TAD callers, further emphasizing the algorithmic challenges and underscoring the need for visual inspection of these results. Finally, we provide links to fully navigable, interactive versions of each of these figures. This gives readers the freedom to explore the full extent of the data outside of the confines of the printed page.

The multiscale nature of Hi-C data demands visualization at a wide range of zoom levels. Its size necessitates piecewise loading of small chunks of data. While genome browsers pioneered multiscale, genome-wide views of 1D data and other tools extended the notion to Hi-C data, the methods of comparison have largely been limited to either a simple vertical tiling of horizontal data tracks or a splitting of Hi-C contact maps along the diagonal. With HiGlass, we have generalized the approach to comparison and extended it beyond simple stacking or two-way splits. We have introduced operations for linking views by location and/or zoom level and for projecting viewports across views. The tool that we have developed, while originally designed for Hi-C data, is a data-agnostic multi-dimensional viewer. Our public demo (http://higlass.io) demonstrates how HiGlass can be used as a standalone viewer to display 1D genomic data [48] while simultaneously providing the same view composition operations for comparison across loci and resolutions.

Having effective tools for comparing genomic data highlights the challenge of organizing such data so that it can be easily found and displayed. Projects such as ENCODE and 4D Nucleome are generating Hi-C data, annotating it with metadata and making it available to the broader public. Efforts like UCSC Genome Browser’s track hubs paved the way for remote genomic data hosting, integration, and visualization. However, there is a need to make it easier for researchers to find and integrate the data that helps answer their biological questions. Future goals in that direction include adding extended metadata to HiGlass data servers and implementing standardized APIs to identify, describe, and query genomic data sets. With more available data, we can take advantage of HiGlass’s extensible architecture to create new ways of exploring, comparing, and interpreting multi-scale experimental results.

## Methods

HiGlass is designed as a client-server application (Figure 6). The client-side user interface is written in JavaScript while the server is written in Python. The client is responsible for arranging tracks and views and requesting data from the server. The server loads data from files in small chunks called “tiles” and sends them back to the client upon request.

### Data are organized according to zoom level using an aggregation or downsampling function

We maintain data at different pre-computed resolutions and when the user zooms in, HiGlass displays higher resolution data. This approach is also employed by web-based map visualization tools such as Google Maps and Open Street Maps. The UCSC Genome Browser and the Integrative Genome Viewer pioneered this approach for genomic data [49,50]. For contact matrices which are generated by binning lists of contacts, creating lower resolution matrices simply requires binning with a larger bin size. The bin sizes used by HiGlass are typically multiples of the powers of 2, starting from the highest resolution data (e.g. for 1K data, bin sizes would be 1K, 2K, 4K, …, 16.384M) but can also be set to arbitrary multiples of the highest resolution. The lower zoom level corresponds to the minimum bin size which can fit 1 / 256th of the width of the matrix. Lower-resolution matrices of counts can also be created by downsampling or “aggregating” higher resolution matrices. In this operation, adjacent pairs of higher resolution bins are merged by summing their values.

For quantitative 1D data, such as RNA-seq or ChIP-seq, the same aggregation procedure can be applied to the 1D array of base-pair resolution values. Adjacent bins are merged by summing their values. In so doing, we maintain a separate array of counts for the number of missing values encountered. This allows us to compute average values when displaying lower resolution data.

For categorical data, downsampling requires discarding values. Values to be discarded are chosen according to an “importance value”. This importance value can be either user-defined or set randomly. A more intelligent importance value can consider a relevant property of the data when deciding which should be visible at lower resolution. For example, for gene annotation tracks, we use a custom importance value based on the number of citations referencing a particular gene. Genes which are well studied and referenced often in the literature, such as TP53 and TNF remain visible as the user zooms out. More obscure genes appear only when there is enough space. For 2D annotations, we use the size of the annotation as an importance value so that larger annotations are visible when zoomed out and smaller annotations only appear at high resolution.

### Tiles break down large datasets into manageable chunks that can be sent from the server to the client

A tile, in the context of HiGlass, is the data available for a given location and zoom level. This is analogous to the tiles used by online maps to show the portion of the map that is visible in the current viewport (Supp. Fig. 7). In the case of Hi-C data, which can be represented as a matrix for any given resolution, a tile consists of a 256 × 256 slice of the matrix.

Zoom levels correspond to the different levels of resolution. The highest zoom level, z_max_, corresponds to the highest resolution data. Each lower zoom level (z-1), corresponds to data at half the resolution of the previous level (r / 2). The data at zoom level 0 must be at a resolution low enough such that the whole genome can be fit into one 256 × 256 tile. This yields an expression for calculating the maximum zoom level for data with a starting (highest) resolution of r_0_ and a genome size of g:

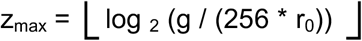

For quantitative 1D genomic data, such as RNA-seq or ChIP-seq or any other coverage-based measure, a tile consists of the data from a 1024 base pair region of genome. The concepts behind the resolution and zoom levels are the same as for 2D data except that instead of a tile corresponding to a square of the matrix at a resolution, it corresponds to a segment of the genome at a given resolution. For qualitative data, the server returns all entries which intersect the length or area of the tile.

In both 1D and 2D data, the lowest resolution is shown at zoom level 0. Given a zoom level, z, the tile visible at genome location *l_g_* can be calculated by considering the width of a tile: *t_w_* = *r*_0_* 2^*z*^

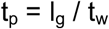

Genomes, being composed of chromosomes, don’t have absolute positions. To get around this, we impose a chromosome ordering for every dataset that is viewable in HiGlass. This must be specified when the data are preprocessed.

### HiGlass stores multi-scale datasets

Due to the limitations of the visible display, there is a fixed amount of data that can be shown in any given area. For a window that is 1024 × 1024 pixels in size, the maximum resolution that the human genome can be shown at is approximately 3 million base pairs / pixels. Fetching all the data from the server is wasteful and unnecessary. We therefore use file formats that store Hi-C and genomic data at multiple resolutions. For Hi-C data, we use the cooler (http://github.com/mirnylab/cooler) format and for genomic data we support the widely used bigWig format [50]. Both support the basic query format of resolution / location. When creating multi-resolution cooler files, we create resolutions that are multiples of the powers of 2 in order to create a smooth transition as the user zooms in and out of the data. While this does increased the size of the data (Supp. Table 2), multiple resolutions are necessary to limit the amount of data that needs to be retrieved from the server when viewing large portions of the contact map.

### The HiGlass server fetches data from files and returns it to the client on demand

The HiGlass server is the interface between the client and the data (Figure 6, Supp. Methods). It receives requests for data (tiles) from the client, opens the data files, and returns only the data requested. This minimizes the amount of data that needs to be sent across the network and in turn lowers the time required to load the data for a given location. Of the 2,333,420 tile requests to our public server at http://higlass.io since February 2017, 2,251,251 (> 96.6%) were fetched with a latency of less 0.5 seconds, a limit beyond which the rates of “observation, generalization and hypothesis significantly decreased” in a controlled user study [51].

The server also maintains a registry of available data files. The client can request a list of available files to provide the user with an overview of data that is available for display. To view data in HiGlass, it first needs to be loaded into the server. Loading the data is done through either a network request or a command line utility.

## Declarations

### Ethics approval and consent to participate

N/A

### Consent for publication

N/A

### Availability of data and material

This paper used data from two published studies, the data for which are on GEO. Note that the data from Rao et al. (Rao et al., 2014) was processed according to the TAD calling procedures described in Forcato et al. (Forcato et al., 2017).

1. Schwarzer et al. (<u>Schwarzer et al. 2017</u>)(Schwarzer et al., 2017): GEO Accession GSE93431 https://www.ncbi.nlm.nih.gov/geo/query/acc.cgi?acc=GSE93431
2. Rao et al. (Rao et al., 2014): GEO Accession GSE63525 https://www.ncbi.nlm.nih.gov/geo/query/acc.cgi?acc=GSE63525

The source code for HiGlass can be found in four complementary repositories

https://github.com/hms-dbmi/higlass - The client side Javascript viewer component
https://github.com/hms-dbmi/higlass-website - A scaffold web site that incorporates the viewer
https://github.com/hms-dbmi/higlass-server - The server we created for serving multi-resolution data
https://github.com/hms-dbmi/higlass-docker - A ready-to-deploy Docker container with installations of the previous three components

Hi-C matrices need to be stored in the cooler format (https://github.com/mirnylab/cooler/) Comprehensive documentation for HiGlass can be found at http://docs.higlass.io

## Funding

This project was made possible by funding from the National Institutes of Health (U01 CA200059, R00 HG007583, and U54 HG007963).

## Author’s Contributions

PK and NG conceived the research. PK, NA, and NG wrote the manuscript with input from LAM, PJP, and BHA. PK, NA, FL, and CM wrote the software with help from KD, HS, JML, SO, AA, NK, JH and SL. BHA, HP, LAM, and PJP provided valuable input and advice for the project.

## Acknowledgements

We thank Francois Spitz, Wibke Schwarzer, Aleksandra Pekowska, Mattia Forcato and Francesco Ferrari for providing the data presented in this paper. We thank Geoffrey Fudenberg for feedback on the manuscript. We also acknowledge important suggestions and feedback from members of the Park Lab at Harvard Medical School, the Mirny Lab at MIT, and the Dekker Lab at University of Massachusetts Medical School as well as members of 4D Nucleome Data Coordination and Integration Center who provided input and feedback. Author Contributions

